# Distinct features of brain perivascular fibroblasts and mural cells revealed by *in vivo* two-photon imaging

**DOI:** 10.1101/2021.05.14.444194

**Authors:** Stephanie K. Bonney, Liam T. Sullivan, Timothy J. Cherry, Richard Daneman, Andy Y. Shih

**Author notes:** **Correspondence:** Andy Y. Shih, Center for Developmental Biology and Regenerative Medicine, Seattle Children’s Research Institute, 1900 9^th^ Ave. M/S JMB-5, Seattle, WA 98101.

## Abstract

Perivascular fibroblasts (PVFs) are recognized for their pro-fibrotic role in many central nervous system disorders. Like mural cells, PVFs surround blood vessels and express Pdgfrβ. However, these shared attributes hinder the ability to distinguish PVFs from mural cells. We used *in vivo* two-photon imaging and transgenic mice with PVF-targeting promoters (Col1a1 or Col1a2) to compare the structure and distribution of PVFs and mural cells in cerebral cortex of healthy, adult mice. We show that PVFs localize to all cortical penetrating arterioles and their pre-capillary offshoots, as well as the main trunk of only larger ascending venules. However, the capillary zone is devoid of PVF coverage. PVFs display short-range mobility along the vessel wall and exhibit distinct structural features (flattened somata and thin ruffled processes) not seen with smooth muscle cells or pericytes. These findings clarify that PVFs and mural cells are distinct cell types coexisting in a similar perivascular niche.

## INTRODUCTION

The microvasculature of the central nervous system (CNS) is surrounded by mural cells and perivascular fibroblasts (PVFs). The physiological roles of these perivascular cell types, and their distinct functions across microvascular zones, continue to be refined with new advances in genetics and imaging^1,2^. Mural cells include vascular smooth muscle cells (SMCs) and different forms of pericytes. SMCs cover surface and penetrating arterioles, and through contractile and dilatory actions of their circumferential cellular processes, dynamically regulate blood flow^3^. Pericytes reside on downstream pre-capillary arterioles and capillaries, and exhibit protruding cell bodies and longer processes with varying degrees of morphological complexity. Ensheathing pericytes on pre-capillary arterioles dynamically regulate blood flow similar to SMCs^4^, while capillary pericytes are involved in slower modulation of capillary tone^5-7^ and blood-brain barrier establishment and homeostasis^8-11^. Unlike mural cells, which are embedded in the endothelial basement membrane^1^, PVFs reside in the perivascular Virchow-Robin space between the mural cell layer and astrocytic endfeet^12,13^. Recent transcriptomic analyses demonstrate that PVFs express genes encoding structural components, modifiers and receptors for the extracellular matrix (ECM)^12,14^. Further, studies in zebrafish have demonstrated that PVFs establish the ECM along developing blood vessels^2^.

PVFs may be a continuation of the pial fibroblast layer of the meninges along blood vessels diving into the neural parenchyma^13,15^. As such, PVFs appear to be most abundant around large diameter arterioles and venules^12,13,16^, however their precise organization along the microvascular tree remains unclear. PVFs also express Pdgfrβ and CD13, both of which are commonly used as broader markers to identify mural cells^16^. This has made it difficult to differentiate PVFs from mural cells and can obscure the distinct physiological roles of these cell types during vascular health and pathology. For example, prior studies have reported that PVFs and meningeal fibroblasts, not mural cells, create the fibrotic scar following CNS injury^14-18^. However, some reports have implicated “type-A pericytes” in this pro-fibrotic response^19,20^ demonstrating the importance of a side-by-side comparison of PVFs to mural cells. Further, since some markers of PVFs are secreted ECM components (collagens), it can be difficult to differentiate PVFs from their secreted proteins by immunostaining. Detailed studies of mural cells in relation to their vascular topology, morphology and dynamics has helped to reveal their unique functions^4,5,21,22^, and now similar analyses are required before PVFs can be carefully studied.

*In vivo* two-photon imaging is an essential approach in the study of neurovascular physiology and pathophysiology. However, it relies on well-characterized transgenic mouse lines to label defined cell types of the neurovascular unit in the healthy brain prior to adding the complexity of tissue reactions to disease and injury. Considering how little is known about PVF distribution and appearance *in vivo*, we leveraged mouse lines with enriched genetic targeting for CNS fibroblasts (Col1a1-GFP and Col1a2-CreERT2) to characterize PVFs in the widely studied vascular architecture of the mouse cerebral cortex. *In vivo* topological analysis revealed that PVFs occupied cortical penetrating arterioles, pre-capillary arteriole and ascending venules, but were absent in capillaries occupied by capillary pericytes across various CNS regions. We further demonstrate that PVFs are morphologically distinct from mural cells, and exhibit short-range mobility on the order of days, in contrast to the stability of mural cells of the adult microvasculature.

## METHODS

### Animals

Mice were housed in specific-pathogen-free facilities approved by AALAC and were handled in accordance with protocols approved by the Seattle Children’s Research Institute IACUC committee. All data were analyzed and reported according to ARRIVE guidelines.

To generate PVF-mural cell reporter mice (Col1a1-GFP; PdgfrβCre-tdTomato), female *Col1a1-GFP/+* mice (C57BL/6 background) were bred with male *Pdgfrβ-Cre/+*; *Ai14-flox/flox* mice. Pdgfrβ-Cre mice (FVB and C57BL/6 × 129 background) were a generous gift from Prof. Volkhard Lindner of the Maine Medical Center Research Institute and the Rosa-lsl-tdTomato reporter mice, Ai14-flox, were obtained from Jackson Labs (#007914; C57BL/6 background). Col1a2-CreER mice (Jackson Lab; #029567; C57BL/6 background) were also crossed with the Ai14-flox and mT/mG-flox lines (#007576) to create tamoxifen inducible PVF-reporter mice. Sparse labeling of PVFs was achieved with two consecutive days of tamoxifen treatment (80mg/kg i.p. dissolved in corn oil). Five consecutive days of tamoxifen treatment (80mg/kg i.p. dissolved in corn oil) resulted in recombination in PVFs and some smooth muscle cells (Supplemental Fig. 6). Mice throughout all experiments were within 3-7 months of age and both males and females were used.

### Cranial window surgery and in vivo two-photon imaging

We created chronic, skull-removed cranial windows over the somatosensory cortex for *in vivo* imaging, as previously described^5^. Briefly, a 3mm in diameter craniotomy was carefully performed with frequently soaking of artificial-cerebral spinal fluid to separate the dura prior to skull removal. After placement and sealing of a 3mm/4mm coverslip plug^7^ and the remaining area surrounding the craniotomy with dental cement, mice were allowed to rest and recover for at least 3 weeks prior to imaging. To label the vasculature in PVF-mural cell reporter mice, 25µL of 5% (w/v in saline) custom conjugated Alexa Fluor 680^23^ (Life Technologies; A20008) conjugated to 2MDa Dextran (Fisher Scientific; NC1275021) was injected through the retro-orbital vein under deep isoflurane anesthesia (2% MAC in medical air). Similarly, the vasculature in PdgfrβCre-tdTomato and Col1a2CreER-tdTomato mice was visualized with 25µL of 5% (w/v in saline) 70kDa FITC-dextran (Sigma-Aldrich; 46945). During imaging, isoflurane was maintained ∼1.5% MAC in medical-grade air. The cortical microvasculature was imaged with a Bruker Investigator coupled to a Spectra-Physics Insight X3. Imaging of Col1a1-GFP; PdgfrβCre-tdTomato and Col1a2CreER-tdTomato mice was performed with 975nm excitation wavelength and 920nm for Col1a1-GFP mice. Collection of green, red and far-red fluorescence emission of PVF-mural cell reporter mice was achieved with 525/70, 595/50, and 660/40 emission bandpass filters respectively, and detected with GaAsP photomultiplier tubes. A 20x (1.0 NA) water-immersion objective (Olympus; XLUMPLFLN) was used to collect high-resolution image z-stacks at 1.0 μm increments. Collection of most images began at the pial surface as identified by pial vessels with penetrating vessels into the cortex. In a subset of image collections, the dural layer was included and recognized by the non-penetrating dural vessels, allowing us to demarcate the meningeal layers in the fibroblast reporter animals.

### Topological analysis of PVFs along the cortical vasculature

Blood flow directionality and vascular structure (in Col1a1-GFP mice) and/or mural cell morphology (in Col1a1-GFP; PdgfrβCre-tdTomato mice) were used to identify penetrating arterioles and ascending venules during *in vivo* two-photon imaging experiments. In total, 21 arteriolar-pre-capillary zones and 32 ascending venules were examined over 4 mice. Branching order off of penetrating arterioles was assigned as described previously^22^. Ascending venules were denoted at 0^th^ order with their respective post-capillary offshoots designated as 1^st^ order. The number of Col1a1-GFP^+^ PVF soma and termination points was documented along branch order. Summation of PVF soma and termination points at their respective branch point was divided by the total number of PVFs or termination points respectively. In all, 80 PVFs and 21 termination points were observed within the arteriolar network and 19 PVFs along ascending venules. FIJI software was used to perform this analysis.

### Penetrating arteriole and ascending venule diameter analysis

Diameter analysis of penetrating arterioles and ascending venules in PVF and PVF-mural reporter mice was performed using the Vasometrics FIJI plug-in^24^. In brief, max projected images were created of penetrating and ascending vessels that included the vessel just under the surface of the brain to their first branch point. The line segment tool was used to draw a through-line along the center of the vessel segment wherein multiple evenly spaced cross-sectional lines were created. The fluorescent intensity profile was created at each cross-section enabling a full-width half max lumen diameter to be calculated along the entire vessel segment. In a few cases, measurement of the penetrating vessel was measured at the pial surface due to the perpendicular nature of certain vessels. Analysis was performed on 15 penetrating arterioles and 32 ascending venules.

### Analysis of perivascular somata roundness

Somata roundness was analyzed using the ImageJ Shape Descriptors measurement tool. Manual measurements outlining perivascular cells were made from maximum projected two-photon images obtained from Col1a2CreER-tdTomato and PdgfrβCre-tdTomato mice. Analysis was performed on 122 perivascular cells along the pre-capillary zone from 8 PdgfrβCre-tdTomato mice and 77 PVFs on penetrating arterioles, pre-capillary zones, and ascending venules from 4 Col1a2CreER-tdTomato mice. To generate a histogram, all data was binned at 0.2 increments and distributed across a scale of roundness where 1 is considered a perfect circle and 0 is completely flat.

### Analysis of PVF and ensheathing pericyte dynamics

Dynamics of PVF and ensheathing pericyte somata was performed on *in vivo* two-photon imaging stacks from Col1a2CreER-tdTomato and PdgfrβCre-tdTomato mice respectively. Distance from the center of individual PVF or ensheathing pericyte soma was measured manually to the nearest capillary junction in a 2D max projected image using the line selection tool in ImageJ. This value and the respective z-distance was then used to calculate the true physical, Euclidean distance at each time point. Displacement over time was calculated by subtracting the Euclidean distance at each time point from their initial Euclidean distance. Analysis was performed on 102 PVF somata from 4 Col1a2CreER-tdtTomato mice and 32 ensheathing pericytes from 5 Pdgfrβ-tdTomato mice.

### Immunohistochemistry and confocal imaging

Mice were deeply anesthetized with euthasol and trans-cardial perfusions were performed with PBS followed by 4% paraformaldehyde. Brains and spinal cord were dissected, cryoprotected in 30% sucrose with 0.001% sodium azide for 1-2 days, frozen in OCT and cryosectioned using a Leica cryostat. To enhance GFP detection, tissue sections (100μm) from Col1a1GFP; PdgfrβCre-tdTomato and Col1a2CreER-mGFP mice animals were stained with rabbit anti-Alexa-fluor 488 conjugated GFP (1:100; ThermoFisher A-21311) and for 48 hours at 4°C followed by 15 minutes of DAPI staining (1:5000; ThermoFisher). Retina whole mounts were also incubated with rabbit anti-Alexa-fluor 488 conjugated GFP for 4 days at 4°C followed by 15 minutes of DAPI staining. Col1a2CreER-tdTomato brain sections (50μm) were stained with rabbit anti-Laminin (1:50; Sigma L9393) or rabbit anti-Pdgfrα (1:100; Cell signaling; D1E1E) overnight at 4°C. Following incubation with these primary antibodies, sections were incubated with appropriate Alexa-fluor conjugated goat anti-rabbit secondary antibodies (1:500; ThermoFisher; A-11008) for 2 hours at room temp followed by 15 minutes of DAPI staining. All antibody staining was performed in a solution of 2% TritonX-100, 10% goat serum and 0.1% sodium azide in PBS. Tissue sections were washed 3 times for 5 minutes in PBS after each antibody incubation. Following staining, tissue was mounted onto slides with Fluoromount G (ThermoFisher; 00-4958-02). Immunofluorescent images were captured using a Leica SP5 or Zeiss 710 LSM confocal microscopes.

### Statistics

All statistical analyses were performed in Graphpad Prism software (ver. 9). Respective statistical analyses are reported in the figure legends. Normality tests, generally Shapiro-Wilk tests, were performed on necessary data sets prior to statistical tests.

## RESULTS

### Perivascular fibroblasts occupy the arteriole, pre-capillary and venule zones but not the capillary bed

To study PVFs and mural cells along the cortical vasculature we generated PVF-mural cell reporter mice (Col1a1-GFP; PdgfrβCre-tdTomato). Both PVFs and mural cells express Pdgfrβ^16^ and therefore are both tdTomato-positive in these mice. However, Col1a1 expression is specific to meningeal fibroblasts (**Supplemental Fig. 1**) and PVFs of the pial surface and intraparenchymal vasculature (**Fig. 1**)^14-17^. Therefore, we were able to identify PVFs in the brain by their additional expression of GFP (**Fig. 1A**).

**Figure 1:**
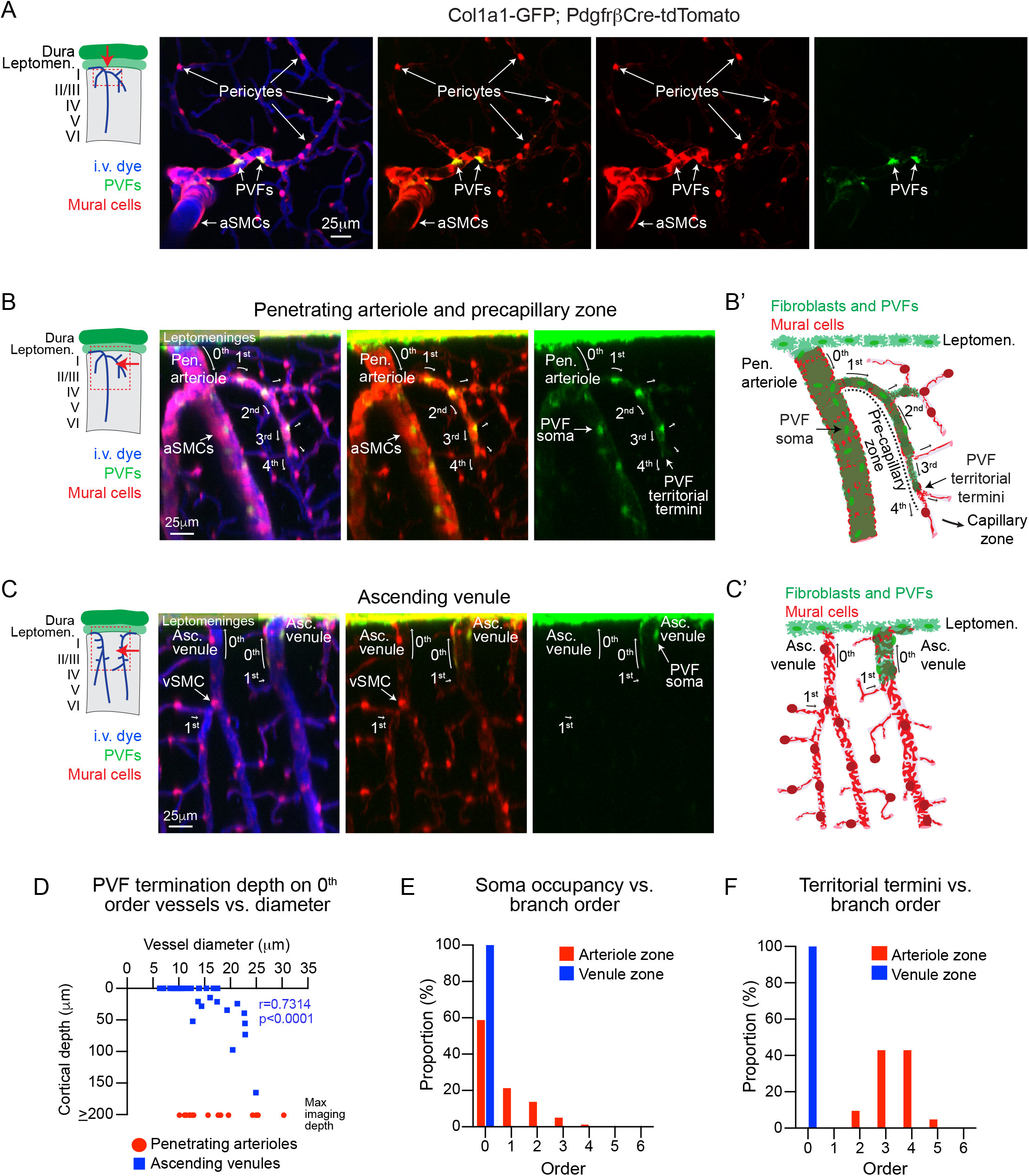
Col1a1-expressing perivascular fibroblasts coexist with mural cells on penetrating arterioles, pre-capillary zones and large ascending venules. **(A)** Top down, max projected image of the cortical vasculature from *in vivo* two-photon imaging of Col1a1-GFP; PdgfrβCre-tdTomato mice showing that tdTomato^+^ mural cells (red) and tdTomato^+^/GFP^+^ perivascular fibroblasts (PVFs; red and green) coexist along the vasculature. Blood vessels labeled with i.v. administration of Alexa-680 (2MDa) (blue). **(B)** Side projection of *in vivo* image stack with (**B’**) schematic demonstrating the topological organization of PVFs (green) along the penetrating arteriole (0^th^ order) as identified by smooth muscle cells (SMCs; red) and pre-capillary zone (1^st^-4^th^ branch order). PVF somata and termination of territory end before the capillary zone (≥ 5^th^ order). **(C)** Side projection of *in vivo* image stack with (**C’**) schematic demonstrating the topological organization of PVFs (green) along the ascending venules (0^th^ order) as identified by venule stellate mural cells (VSMCs; red) and 1^st^ order post-capillary connections. PVF somata only occupy the main trunk (0^th^ order) of some ascending venules. **(D)** Scatter plot of penetrating arteriole (red) and ascending venule (blue) diameters versus PVF termination depth along 0^th^ order vessels. Spearman’s rank correlation shows a positive correlation along ascending venules (****p<0.0001, r=0.7314, n=32 ascending venules). PVFs were observed up to maximum imaging depths along penetrating arterioles regardless of vessel diameter (n=15 penetrating arterioles). Data was compiled from 4 mural cell-PVF and PVF reporter (Col1a1-GFP) mice. **(E)** Histogram depicting the proportion of PVF soma occupancy from 0^th^ to 6^th^ branch order along the arteriole (red) or venule (blue) zones. **(F)** Histogram depicting the proportion of PVF territorial termination from 0^th^ to 6^th^ branch order along the arteriole (red) or venule (blue) zones.

We first used *in vivo* two-photon imaging to assess the organization of PVFs along vascular zones previously defined by hierarchical organization of mural cells within the cortex^7,22^. Penetrating arterioles, denoted as 0^th^ order, were identified by robust tdTomato-expressing SMCs. Pre-capillary arterioles (∼1^st^ – 4^th^ order) branch off the penetrating arterioles and ramify into the capillary zone (>5^th^ order), increasing in branch order with each bifurcation. Here, branch orders 1 – 4^th^ order are considered the pre-capillary zone due to α-smooth muscle actin (α-SMA)-expressing ensheathing pericytes that tend to cover these vessel segments^4,21,22^. Ensheathing pericytes can be identified by the morphology of their processes, which completely encircled the endothelium and are more elongated compared to SMCs. When pericytes shift their morphology to mesh and thin-strand processes at or before 4^th^ order, this represents entry into the α-SMA-low to negative capillary zone (**Fig. 1B**). We analyzed the distribution of PVF somata and the termination of the territories covered by their processes with respect to branch orders. This revealed that PVF somata were predominantly found along 0^th^ order penetrating arterioles, and up to 4^th^ order branches (**Fig. 1B, E**) with their territories terminating predominantly on 3^rd^ and 4^th^ order vessels (**Fig. 1B, F**). This corresponds to the pre-capillary arteriole zone. PVFs were absent within the capillary zone. Over the cortical depths we could image, PVFs were continually present on penetrating arterioles even up to the maximum imaging depth used (≥ 200μm) (**Fig. 1D**). This was the case regardless of penetrating arteriole diameter.

Ascending venules (0^th^ order on the venule side) drain blood out of the brain and are covered by tdTomato-expressing venule stellate-shaped mural cells (VSMCs)^21,25^. We found that PVF somata were present on some, but not all ascending venules (**Fig. 1C**). Only 37.5% (12/32) of 0^th^ order ascending venules were covered by PVFs, and these were typically ascending venules of larger diameter (>12μm). PVF territories along ascending venules also terminated at shallower depths than penetrating arterioles and the depth of termination correlated with the ascending venule diameter (**Fig. 1D**). Unlike arterioles, PVFs on the venous side did not extend beyond the 0^th^ order vessel with either their somata or processes (**Fig. 1E, F**).

Immunohistochemical analysis allowed us to examine penetrating arteriole and ascending venule labeling beyond cortical depths achieved by *in vivo* imaging. We found that PVF coverage of penetrating arterioles extended down to their deepest terminal branches (**Supplemental Fig. 2**). Interestingly, principal cortical venules (PCVs), which are the largest venules that drain cortical gray matter and underlying white matter^26^, exhibited PVF labeling even down to their branches in the external capsule/corpus callosum (**Supplemental Fig. 2**). Together this demonstrates that PVFs are found along penetrating arterioles, pre-capillary zones and the trunk of large diameter ascending venules, but not along capillaries in the cortex. Thus, PVFs and capillary pericytes occupy distinct vascular zones.

### Topological organization of perivascular fibroblasts is identical across various CNS regions

We next asked if PVFs were organized similarly along the vasculature throughout the CNS by performing confocal microscopy on various CNS regions of PVF-mural cell reporter mice. Similar to what we observed in the cortex (**Fig. 1 and Fig. 2A**), PVFs were found on arterioles, pre-capillary arterioles and large diameter venules, but not capillaries, in the corpus callosum (**Fig. 2B, Supplemental Fig. 2**) and hippocampus (**Fig. 2C**), which are two regions where vascular pathology contributes to neurological disease^27,28^. PVFs in the spinal cord are also organized similarly, a CNS region where fibrotic responses elicited by injury are regularly studied (**Fig. 2D**)^14,16,17^. The same topological organization was also observed in the striatum, thalamus and the cerebellum (**Supplemental Fig. 3A-C**). Interestingly, PVFs were not found along the retinal vasculature (**Supplemental Fig. 3D**). This may be due to the fact that the pia covers the optic nerve but does not extend to the retina^29^ and therefore further supports the idea that PVFs are an extension of the pia^15^.

**Figure 2:**
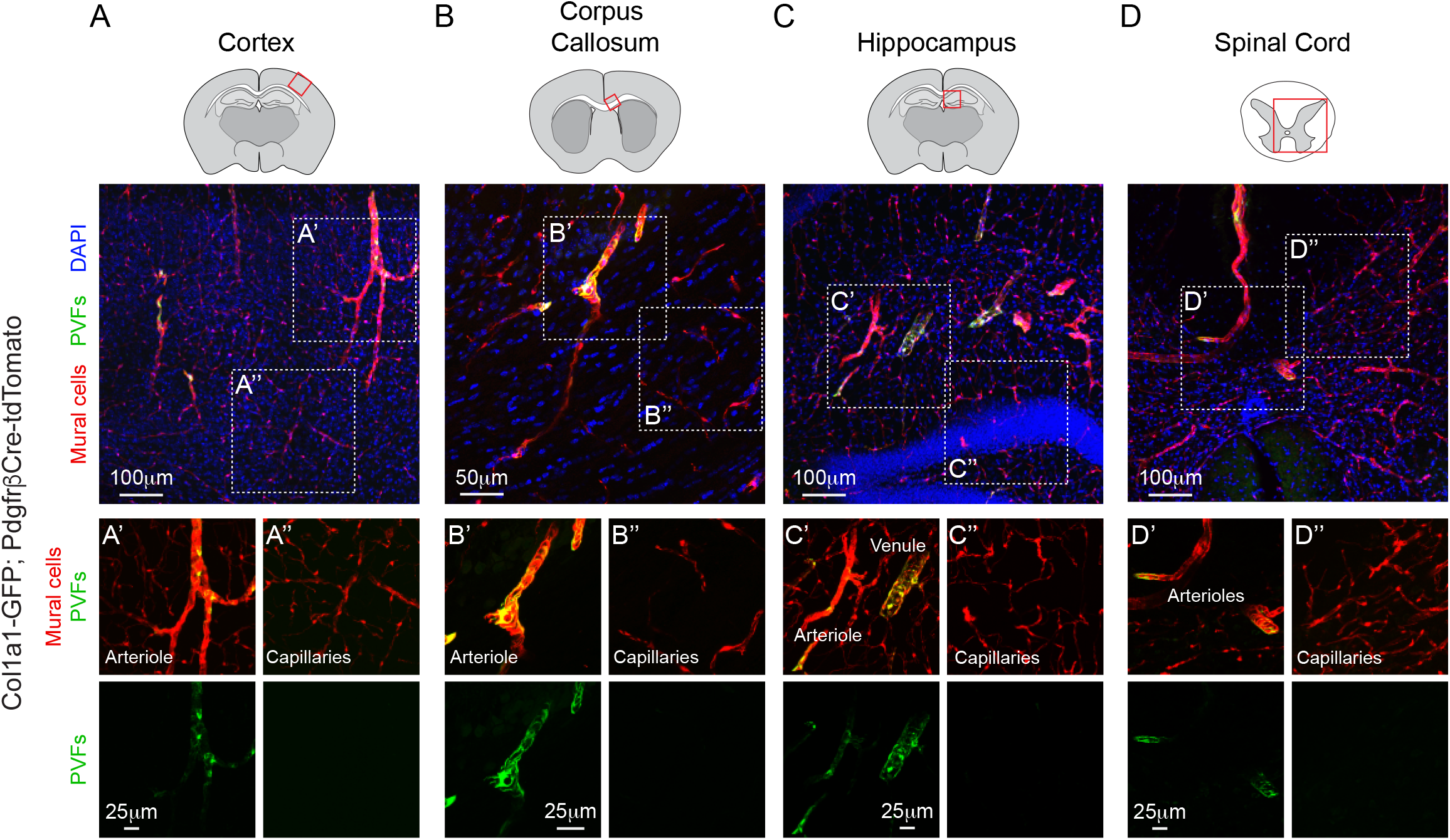
Topological organization of PVFs is similar across CNS regions. Representative confocal image of the **(A)** cortex, **(B)** corpus callosum, **(C)** hippocampus and **(D)** spinal cord from Col1a1-GFP; PdgfrβCre-tdTomato mice. Inset images show that perivascular fibroblasts (PVFs; green) are found along arterioles, pre-capillary zones and venules in the **(A’)** cortex, **(B’)** corpus callosum, **(C’)** hippocampus **(D’)** and spinal cord. However, PVFs are not present on the capillary zone, which is occupied by thin-strand and mesh pericytes (red) in the **(A’’)** cortex, **(B’’)** corpus callosum, **(C’’)** hippocampus **(D’’)** and spinal cord. Mural cells are shown in red and DAPI in blue.

### Morphological features of perivascular fibroblasts are distinct from mural cells

Morphological description of PVFs and a comparison to mural cells would assist in differentiating these two perivascular cell types. The morphology of mural cells along different microvascular zones in cerebral cortex has been described in detail using PdgfrβCre mice^21,25^. Although PVFs are also labeled with the PdgfrβCre driver, as shown above, mural cells are brighter and more conspicuous in PdgfrβCre-tdTomato mice. SMCs along penetrating arterioles have thick circumferentially-oriented processes (**Fig. 3A**). Ensheathing pericytes of the pre-capillary zone also have thick circumferential processes, and exhibit protruding, ovoid cell bodies typical of pericytes (**Fig. 3B**). VSMCs also have protruding cell bodies like pericytes but their processes extend in multiple directions thus take on an overall stellate shape (**Fig. 3C**)^21,25^. Observations from Figure 1 demonstrated flattened PVF somata that did not protrude from the vessel wall. As such, putative PVF somata were occasionally observed along the vasculature of PdgfrβCre-tdTomato mice, most often along the pre-capillary zone where morphological distinction were the greatest (**Fig. 3B and Supplemental Fig. 4**).

**Figure 3:**
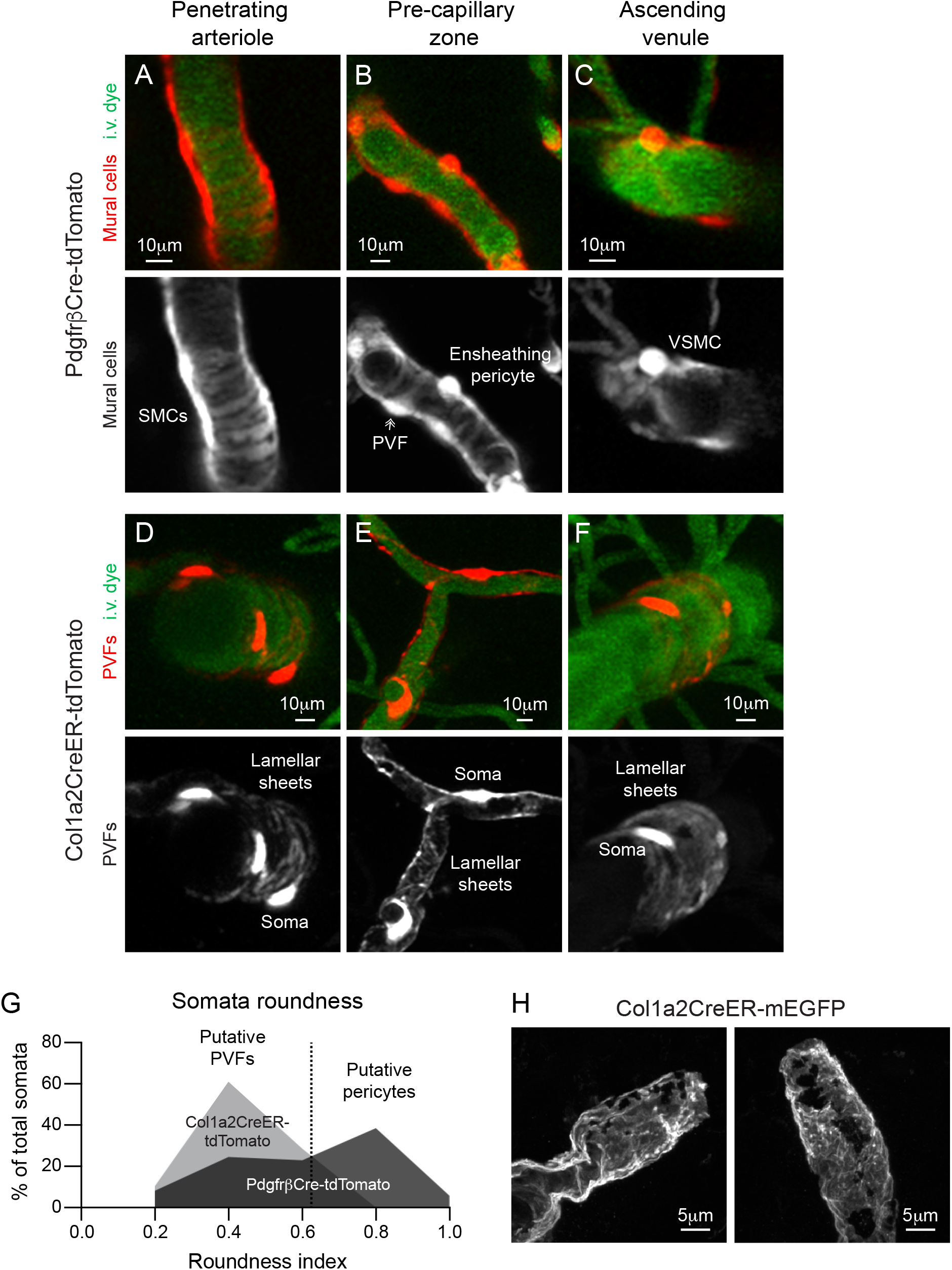
Morphological features of PVFs are distinct from mural cells. **(A-C)** Representative two-photon images from PdgfrβCre-tdTomato mice of **(A)** smooth muscle cells (SMCs) on a penetrating arteriole, **(B)** ensheathing pericyte and a putative PVF in the pre-capillary zone and **(C)** venule stellate mural cells (VSMCs) on an ascending venule with respective red channel separated to better show mural cell morphology (white). Vasculature labeled with i.v. administration of FITC-dextran (70kDa; green). **(D-F)** Representative two-photon images of PVFs from Col1a2CreER-tdTomato mice treated with 2 consecutive days of tamoxifen (80mg/kg) (red) on the **(D)** penetrating arteriole, (**E)** pre-capillary zone and **(F)** ascending venule with respective red channel separated below to appreciate PVF morphology in white. Vasculature labeled with i.v. administration of FITC-dextran (70kDa; green). **(G)** Histogram of somata roundness of tdTomato+ perivascular cells along the pre-capillary zone in PdgfrβCre-tdTomato mice (black) and tdTomato+ PVFs along penetrating arterioles, pre-capillary zone, and ascending venules in Col1a2CreER-tdTomato mice (gray). A roundness index of 1 would be considered a perfect circle. Data was compiled from 122 perivascular cells from 8 PdgfrβCre-tdTomato mice and 77 PVFs from 4 Col1a2CreER-tdTomato mice **(H)** Representative high-resolution confocal image of ruffled PVF membrane (white) from Col1a2CreER-mGFP mice.

To gain a clearer view of PVF morphology, we utilized the tamoxifen inducible fibroblast-targeting line Col1a2CreER and crossed them with the Rosa-lsl-tdTomato reporter mice (Ai14-flox)^30^. Recent work showed that Col1a2CreER cell targeting overlaps with ∼90% of Col1a1-GFP expressing fibroblasts in the CNS^14^. Col1a2CreER-tdTomato mice were given two days of tamoxifen (80mg/kg) to sparsely label PVFs. Two-photon imaging revealed sparse tdTomato expression in PVFs of brain arterioles and venules, with vascular zone specificity identical to GFP in Col1a1-GFP mice. Meningeal fibroblasts were also sparsely labeled (**Supplemental Fig. 5**). Expression of a second fibroblast marker, Pdgfrα, was also confirmed in PVFs along cortical vessels in Col1a2CreER-tdTomato mice. Five days of tamoxifen resulted in tdTomato expression in some SMCs, indicating that tamoxifen doseage is important for precise labeling of PVFs (**Supplemental Fig. 6**). Together, these observations verify that Col1a2CreER can be a specific and efficient Cre driver to study isolated fibroblasts and PVFs in the CNS.

Imaging of PVF-reporter mice revealed that PVFs on penetrating arterioles, pre-capillary zones and ascending venules all shared similar features: Flattened somata and thin, sheet-like lamella that covered much of the vessel surface (**Fig. 3D-F**). Analysis of perivascular somata roundness along the pre-capillary zone in PdgfrβCre-tdTomato mice revealed two groups of somata shapes. One group with a round shape (∼>0.6 roundness index), putative ensheathing pericytes, and another group with a more flattened somata (∼<0.6 roundness index), putative PVFs. Indeed, the roundness index of PVF somata in Col1a2CreER-tdTomato fell between 0.2-0.7, coinciding with the flattened somata group in the PdgfrβCre-tdTomato mice (**Fig. 3G**).

We also examined PVFs in Col1a2CreER-mGFP mice by high resolution confocal microscopy, where cells expressed membrane-bound EGFP to improve visibility of fine subcellular structure. This revealed that PVF processes were fibrous sheets with “ruffled” texture (**Fig. 3H, Supplementary Fig. 7**). This is consistent with electron microscopy studies of human cortical arterioles and venules, which are described to be surrounded by thin “pial cells” within perivascular spaces^13^. Thus, PVFs are morphologically distinct from mural cells both at the level of their somata (protruding vs flattened) and processes (circumferential banding vs. ruffled sheets).

### Perivascular fibroblasts are a dynamic perivascular cell population

Previous studies showed that the somata of pericytes were very stable in their positions^5^. This is in line with their tight physical interlocking with endothelial cells and encasement in vascular basement membrane^31^. In contrast, PVFs reside within the fluid-filled perivascular space of large vessels, raising the possibility that PVFs can move more easily through this space. We therefore monitored the position of PVF somata weekly up to 4 weeks in most cases. To pinpoint their position from week to week we measured the Euclidian distance from the center of an individual PVF soma to the nearest capillary branch point, a vascular structure known to be stable over long periods in adult cortex^32^. PVFs often made small morphological adjustments to their somata overtime (**Fig. 4A**). Some PVFs exhibited clear mobility as their somata displaced from their initial positions on average 5μm and up to 12μm (**Fig. 4B, E**). This movement may not be a passive effect of cerebral spinal fluid (CSF) flow^33^, as PVFs appeared to leave and return to the same locations over time (**Fig. 4B**). Mobile PVFs were typically on locations away from vascular junctions (non-junctional), whereas PVFs located at junctions (junctional) were generally more stable (**Fig. 4C, F**). These aspects of PVF dynamics were similar across the different vascular zones (**Supplemental Fig. 8**). Thus, PVFs are dynamic but structural limitations of the vessel may restrict their movements. This contrasts with ensheathing pericytes, which occupied the same zone as the PVFs examined, but display little, if any movement over 28 days (**Fig. 4D, G**). Composite data across experiments confirmed that PVFs were significantly more mobile than ensheathing pericytes (**Fig. 4H**).

**Figure 4:**
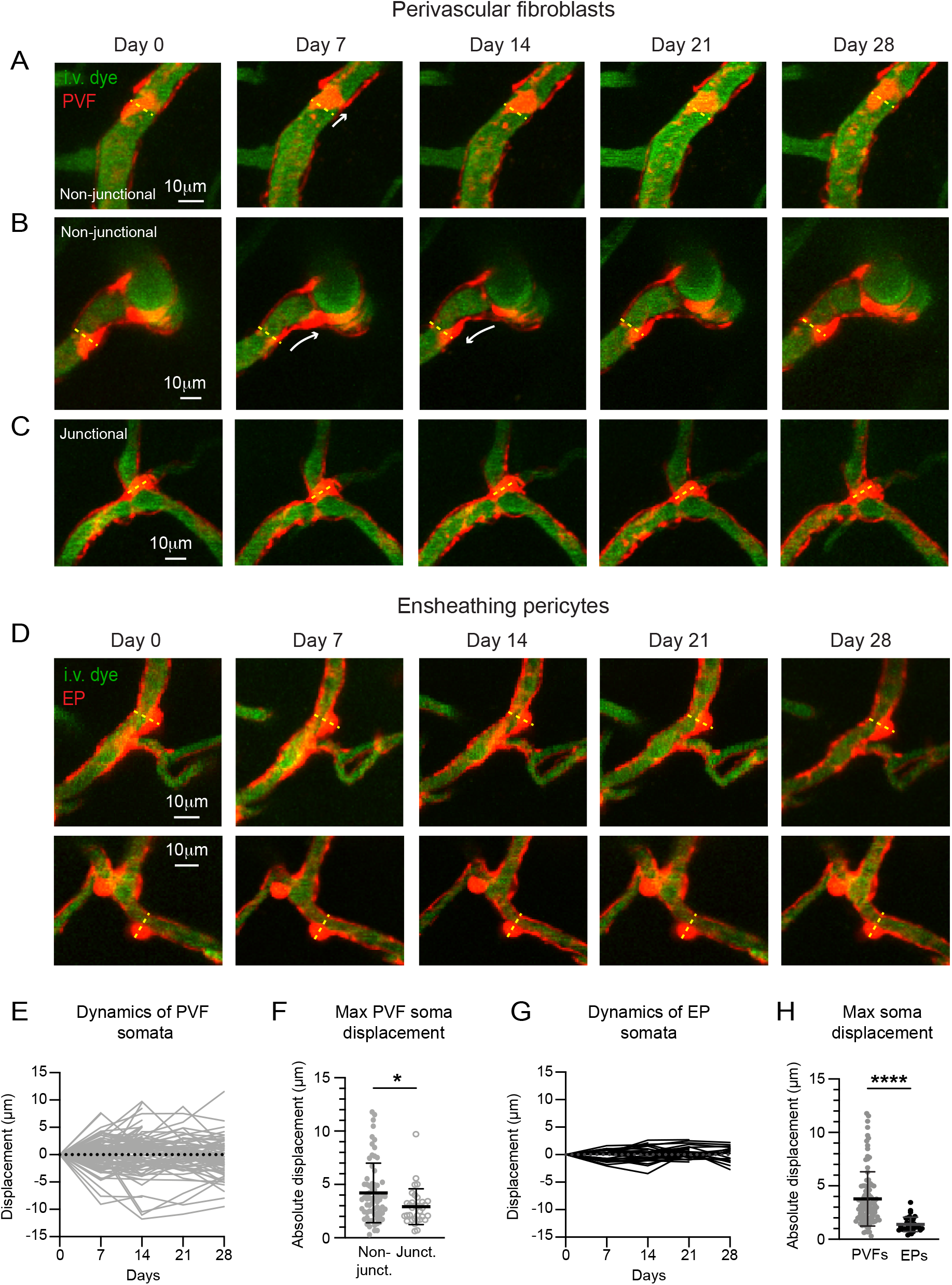
Perivascular fibroblasts are a dynamic perivascular cell population. **(A-C)** Representative *in vivo* two-photon images of perivascular fibroblasts (PVF; red) over 28 days from Col1a2CreER-tdTomato mice. **(A, B)** Non-junctional PVFs display more mobility than **(C)** PVFs found at vascular junctions. Dashed line indicates initial soma position on Day 0. Vasculature labeled with i.v. administration of FITC-dextran (70kDa; green). **(D)** Two representative examples of ensheathing pericytes from PdgfrβCre-tdTomato mice following 28 days of *in vivo* two-photon imaging. Dashed line indicates initial soma position on Day 0. Vasculature labeled with i.v. administration of FITC-dextran (70kDa; green). **(E)** Graph demonstrating soma displacement of PVFs over 28 days from initial position on day 0 (n=102 PVFs from 4 Col1a2CreER-tdTomato mice). **(F)** Graph comparing maximum displacement of PVFs with non-junctional (n=68 PVFs) and junctional (n=34) vascular positions. PVFs with non-junctional positions were significantly more dynamic than PVFs at junctions (Mann-Whitney test U=841, *p=0.0251. Median: Non-junctional=3.627, junctional=2.814). **(E)** Graph demonstrating soma displacement of ensheathing pericytes (EP) over 28 days from initial position on day 0 (n=32 EPs from 5 PdgfrβCre-tdTomato mice). **(H)** Graph comparing maximum displacement of PVFs (n=102 PVFs) and EPs (n=32). PVFs were significantly more dynamic than EPs (Mann-Whitney test U=431, ****p<0.0001. Median: PVFs=3.081, EP=1.142).

## DISCUSSION

Our studies confirm that PVFs and mural cells co-exist in a similar perivascular niche in brain microvasculature, but cell-specific transgenic mouse lines and morphological distinctions can be used to differentiate the cell types. The data shown here provides foundational knowledge on PVF organization, morphology and dynamics that will be necessary before detailed studies on cellular reactions to disease or injury using *in vivo* two-photon microscopy. Future studies will be needed to delineate the physiological role of PVFs in the healthy brain. For example, by residing solely on arterioles, pre-capillary zones and larger ascending venules PVFs may play a unique role in large vessel stability. The parenchymal basement membrane outside of arterioles is a fluid filled perivascular space occupied by PVFs intermixed with a composition of laminin-α1/2 and fibrillar Col1 and Col3^34-36^. In addition to Col1 expression, transcriptional analysis suggests that PVFs also express Col3 and laminin-α1/2 and may therefore be the source of these ECM components^12,14^. In zebrafish, deposition of collagen (Col1a2 and Col5a1) by PVFs was vital for vascular stability along intersegmental vessels that sprout off of the dorsal aorta^2^. Interestingly, upon spreading out their fibrous lamella, fibroblasts form actin stress fibers and focal adhesions attaching to the ECM and help create tension within their environment^37^. Thus, the sheet-like encasement of PVF processes may physically strengthen the vascular basement membrane. This possibility could be tested by optical ablation of resting PVFs along the brain vasculature or deletion of PVF-derived basement membrane proteins.

We also demonstrate that PVFs are dynamic along the brain vasculature. It’s possible that PVFs are actively responding to environmental cues or migratory signals during their movement. The small adjustments seen in PVF position could also be a passive effect of vasomotion, the natural slow oscillations (∼0.1Hz) in arteriole diameter important for driving CSF flow^33^. Although, we did not observe net directional movement of PVFs, as might be expected if PVFs migrated with flow of CSF. Vasomotion is substantially reduced under isoflurane-induced anesthesia, as used in these studies^38^, and therefore future experiments could assess this possibility in awake mice. We were unable to resolve distinct territories of individual PVFs in instances where multiple tdTomato-expressing PVFs were present along the vasculature. It is possible they overlap with one another, and this was a limitation in assessing the dynamics of PVF lamellar sheets.

Vascular stability and clearance of brain waste through perivascular spaces is substantially altered in aging and Alzheimer’s related diseases^39-41^. Native PVF morphology can change with their environmental conditions. For example, fibroblasts in high tension environments, such as those created by Col1 and 3, spread out their membrane and form lamellar structures as opposed to dendritic morphologies exhibited by fibroblasts in low-tension environments^42,43^. Thus, under conditions of small vessel disease or tissue hypoxia, PVFs could conceivably have an effect on glymphatic drainage by obstructing or changing the shape of the perivascular space along arterioles and venules. Indeed, activation of PVFs coincided with enlarged perivascular spaces in patient tissue and mouse models of amyotrophic lateral sclerosis^44^. Further, if PVFs do contribute to the composition of the vascular basement membrane, PVF pathology may alter the intramural peri-arterial drainage for CSF^45^. Both of these CSF drainage pathways have been implicated in removal of waste metabolites and amyloid beta from the brain. These unexplored mechanisms within the perivascular space would be in addition to the overt migration of PVFs from vessels to form a component of tissue scars following injury, which has thus far been the main focus of PVFs in disease.

## ACKNOWLEDGMENTS

This work was support by NIH NRSA fellowship to SKB F32 NS117649 and grants to AYS (NS106138, AG063031, NS097775). The authors greatly appreciate the fruitful discussions from members of the Shih Lab, specifically Drs. Andrée-Anne Berthiaume, Vanessa Coelho-Santos and Stefan Stamenkovic. We are also thankful for the discussions with Drs. Julie Siegenthaler and Mark Majesky throughout these studies.

## AUTHOR CONTRIBUTIONS

SKB and AYS conceptualized and designed experiments. Two-photon imaging and data analysis was done by SKB. Tissue collection and confocal imaging was performed by LTS and SKB. Retina dissections were assisted by TJC. Statistics was performed by SKB. Manuscript was written by SKB and AYS with editing and contributions from all authors.

## REFERENCES

1. Sweeney MD, Ayyadurai S, Zlokovic BV. Pericytes of the neurovascular unit: key functions and signaling pathways. Nat Neurosci. 2016;19(6):771–783.

2. Rajan AM, Ma RC, Kocha KM, Zhang DJ, Huang P. Dual function of perivascular fibroblasts in vascular stabilization in zebrafish. PLoS Genet. 2020;16(10):e1008800.

3. Kisler K, Nelson AR, Montagne A, Zlokovic BV. Cerebral blood flow regulation and neurovascular dysfunction in Alzheimer disease. Nat Rev Neurosci. 2017;18(7):419–434.

4. Gonzales AL, Klug NR, Moshkforoush A, et al. Contractile pericytes determine the direction of blood flow at capillary junctions. Proc Natl Acad Sci U S A. 2020;117(43):27022–27033.

5. Berthiaume AA, Grant RI, McDowell KP, et al. Dynamic Remodeling of Pericytes In Vivo Maintains Capillary Coverage in the Adult Mouse Brain. Cell Rep. 2018;22(1):8–16.

6. Hall CN, Reynell C, Gesslein B, et al. Capillary pericytes regulate cerebral blood flow in health and disease. Nature. 2014;508(7494):55–60.

7. Hartmann DA, Berthiaume AA, Grant RI, et al. Brain capillary pericytes exert a substantial but slow influence on blood flow. Nat Neurosci. 2021.

8. Daneman R, Zhou L, Kebede AA, Barres BA. Pericytes are required for blood-brain barrier integrity during embryogenesis. Nature. 2010;468(7323):562–566.

9. Armulik A, Genové G, Mäe M, et al. Pericytes regulate the blood-brain barrier. Nature. 2010;468(7323):557–561.

10. Ben-Zvi A, Lacoste B, Kur E, et al. Mfsd2a is critical for the formation and function of the blood-brain barrier. Nature. 2014;509(7501):507–511.

11. Nikolakopoulou AM, Montagne A, Kisler K, et al. Pericyte loss leads to circulatory failure and pleiotrophin depletion causing neuron loss. Nat Neurosci. 2019;22(7):1089–1098.

12. Vanlandewijck M, He L, Mäe MA, et al. A molecular atlas of cell types and zonation in the brain vasculature. Nature. 2018;554(7693):475–480.

13. Zhang ET, Inman CB, Weller RO. Interrelationships of the pia mater and the perivascular (Virchow-Robin) spaces in the human cerebrum. J Anat. 1990;170:111–123.

14. Dorrier CE, Aran D, Haenelt EA, et al. CNS fibroblasts form a fibrotic scar in response to immune cell infiltration. Nat Neurosci. 2021;24(2):234–244.

15. Kelly KK, MacPherson AM, Grewal H, et al. Col1a1+ perivascular cells in the brain are a source of retinoic acid following stroke. BMC Neurosci. 2016;17(1):49.

16. Soderblom C, Luo X, Blumenthal E, et al. Perivascular fibroblasts form the fibrotic scar after contusive spinal cord injury. J Neurosci. 2013;33(34):13882–13887.

17. Yahn SL, Li J, Goo I, Gao H, Brambilla R, Lee JK. Fibrotic scar after experimental autoimmune encephalomyelitis inhibits oligodendrocyte differentiation. Neurobiol Dis. 2020;134:104674.

18. Fernández-Klett F, Potas JR, Hilpert D, et al. Early loss of pericytes and perivascular stromal cell-induced scar formation after stroke. J Cereb Blood Flow Metab. 2013;33(3):428–439.

19. Göritz C, Dias DO, Tomilin N, Barbacid M, Shupliakov O, Frisén J. A pericyte origin of spinal cord scar tissue. Science. 2011;333(6039):238–242.

20. Dias DO, Kim H, Holl D, et al. Reducing Pericyte-Derived Scarring Promotes Recovery after Spinal Cord Injury. Cell. 2018;173(1):153-165.e122.

21. Hartmann DA, Underly RG, Grant RI, Watson AN, Lindner V, Shih AY. Pericyte structure and distribution in the cerebral cortex revealed by high-resolution imaging of transgenic mice. Neurophotonics. 2015;2(4):041402.

22. Grant RI, Hartmann DA, Underly RG, Berthiaume AA, Bhat NR, Shih AY. Organizational hierarchy and structural diversity of microvascular pericytes in adult mouse cortex. J Cereb Blood Flow Metab. 2019;39(3):411–425.

23. Li B, Ohtomo R, Thunemann M, et al. Two-photon microscopic imaging of capillary red blood cell flux in mouse brain reveals vulnerability of cerebral white matter to hypoperfusion. J Cereb Blood Flow Metab. 2020;40(3):501–512.

24. McDowell KP, Berthiaume AA, Tieu T, Hartmann DA, Shih AY. VasoMetrics: unbiased spatiotemporal analysis of microvascular diameter in multi-photon imaging applications. Quant Imaging Med Surg. 2021;11(3):969–982.

25. Hill RA, Tong L, Yuan P, Murikinati S, Gupta S, Grutzendler J. Regional Blood Flow in the Normal and Ischemic Brain Is Controlled by Arteriolar Smooth Muscle Cell Contractility and Not by Capillary Pericytes. Neuron. 2015;87(1):95–110.

26. Duvernoy HM, Delon S, Vannson JL. Cortical blood vessels of the human brain. Brain Res Bull. 1981;7(5):519–579.

27. Montagne A, Nikolakopoulou AM, Zhao Z, et al. Pericyte degeneration causes white matter dysfunction in the mouse central nervous system. Nat Med. 2018;24(3):326–337.

28. Uemura MT, Maki T, Ihara M, Lee VMY, Trojanowski JQ. Brain Microvascular Pericytes in Vascular Cognitive Impairment and Dementia. Front Aging Neurosci. 2020;12:80.

29. Kur J, Newman EA, Chan-Ling T. Cellular and physiological mechanisms underlying blood flow regulation in the retina and choroid in health and disease. Prog Retin Eye Res. 2012;31(5):377–406.

30. Zheng B, Zhang Z, Black CM, de Crombrugghe B, Denton CP. Ligand-dependent genetic recombination in fibroblasts : a potentially powerful technique for investigating gene function in fibrosis. Am J Pathol. 2002;160(5):1609–1617.

31. Ornelas S BA, Bonney SK, Coelho-Santos V, Underly RG, Kremer A, Guerin CJ, Lippens S, and Shih AY. Three-dimensional ultrastructure of the brain pericyte-endothelial interface. In. (In press) ed: JCBFM; 2021.

32. Harb R, Whiteus C, Freitas C, Grutzendler J. In vivo imaging of cerebral microvascular plasticity from birth to death. J Cereb Blood Flow Metab. 2013;33(1):146–156.

33. Iliff JJ, Wang M, Zeppenfeld DM, et al. Cerebral arterial pulsation drives paravascular CSF-interstitial fluid exchange in the murine brain. J Neurosci. 2013;33(46):18190–18199.

34. Lam MA, Hemley SJ, Najafi E, Vella NGF, Bilston LE, Stoodley MA. The ultrastructure of spinal cord perivascular spaces: Implications for the circulation of cerebrospinal fluid. Sci Rep. 2017;7(1):12924.

35. Hannocks MJ, Pizzo ME, Huppert J, et al. Molecular characterization of perivascular drainage pathways in the murine brain. J Cereb Blood Flow Metab. 2018;38(4):669–686.

36. Thomsen MS, Routhe LJ, Moos T. The vascular basement membrane in the healthy and pathological brain. J Cereb Blood Flow Metab. 2017;37(10):3300–3317.

37. Petroll WM, Ma L, Jester JV. Direct correlation of collagen matrix deformation with focal adhesion dynamics in living corneal fibroblasts. J Cell Sci. 2003;116(Pt 8):1481–1491.

38. van Veluw SJ, Hou SS, Calvo-Rodriguez M, et al. Vasomotion as a Driving Force for Paravascular Clearance in the Awake Mouse Brain. Neuron. 2020;105(3):549-561.e545.

39. Kress BT, Iliff JJ, Xia M, et al. Impairment of paravascular clearance pathways in the aging brain. Ann Neurol. 2014;76(6):845–861.

40. van Veluw SJ, Scherlek AA, Freeze WM, et al. Different microvascular alterations underlie microbleeds and microinfarcts. Ann Neurol. 2019;86(2):279–292.

41. Iliff JJ, Wang M, Liao Y, et al. A paravascular pathway facilitates CSF flow through the brain parenchyma and the clearance of interstitial solutes, including amyloid β. Sci Transl Med. 2012;4(147):147ra111.

42. Rhee S. Fibroblasts in three dimensional matrices: cell migration and matrix remodeling. Exp Mol Med. 2009;41(12):858–865.

43. Xu J, Shi GP. Vascular wall extracellular matrix proteins and vascular diseases. Biochim Biophys Acta. 2014;1842(11):2106–2119.

44. Månberg A, Skene N, Sanders F, et al. Altered perivascular fibroblast activity precedes ALS disease onset. Nat Med. 2021;27(4):640–646.

45. Hawkes CA, Härtig W, Kacza J, et al. Perivascular drainage of solutes is impaired in the ageing mouse brain and in the presence of cerebral amyloid angiopathy. Acta Neuropathol. 2011;121(4):431–443.

