## Supplementary Figures for "Distinct features of brain perivascular fibroblasts and mural cells revealed by *in vivo* two-photon imaging"

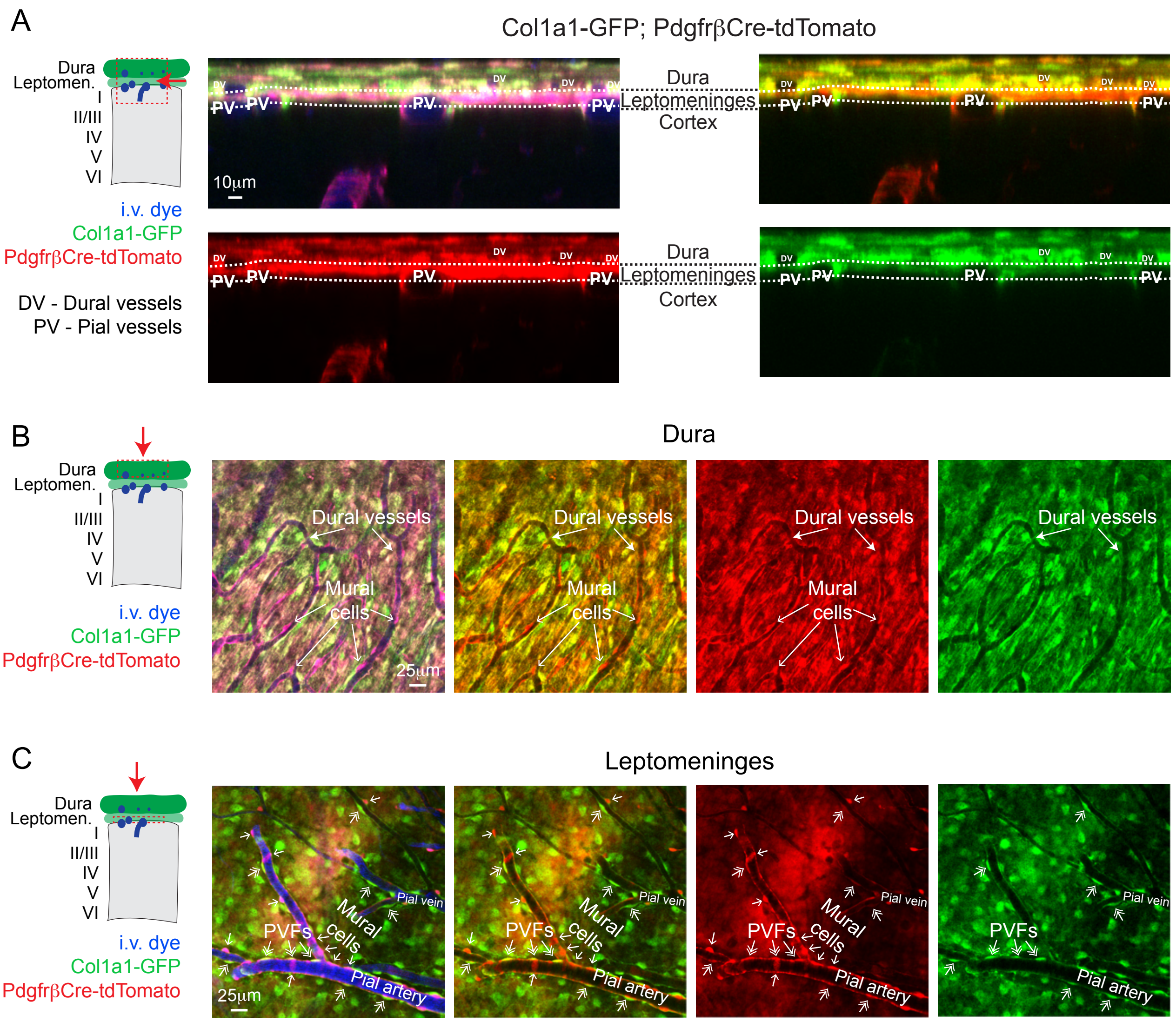


**Supplement Figure 1: Cortical meninges are made up of Col1a1+/Pdgfrβ+ meningeal fibroblasts.**

**(A)** Side projection of the meningeal layers (dura and leptomeninges) and cortex from *in vivo* two-photon imaging of Col1a1-GFP; PdgfrβCre-tdTomato mice showing shared expression of PdgfrβCre-tdTomato (red) and Col1a1-GFP (green) in both layers. Vasculature labeled with i.v. administration of Alexa-680 dextran (2MDa) (blue). Presence of dural vessels (DV) and pial vessels (PV) allowed for the differentiation of the different meningeal layers.
**(B)** Top down, max projection of the dural layer demonstrated that Col1a1+/Pdgfrβ+ fibroblasts make up the dura. Pdgfrβ+ mural cells (arrows; tdTomato+) were also present on dural vessels.
**(C)** Top down, max projection of leptomeninges showing these layers (arachnoid and pia) were made up of Col1a1+/Pdgfrβ+ fibroblasts. Perivascular fibroblasts (PVFs) (double arrows; GFP+/tdTomato+) and mural cells (arrows; tdTomato+) are present on pial vessels.

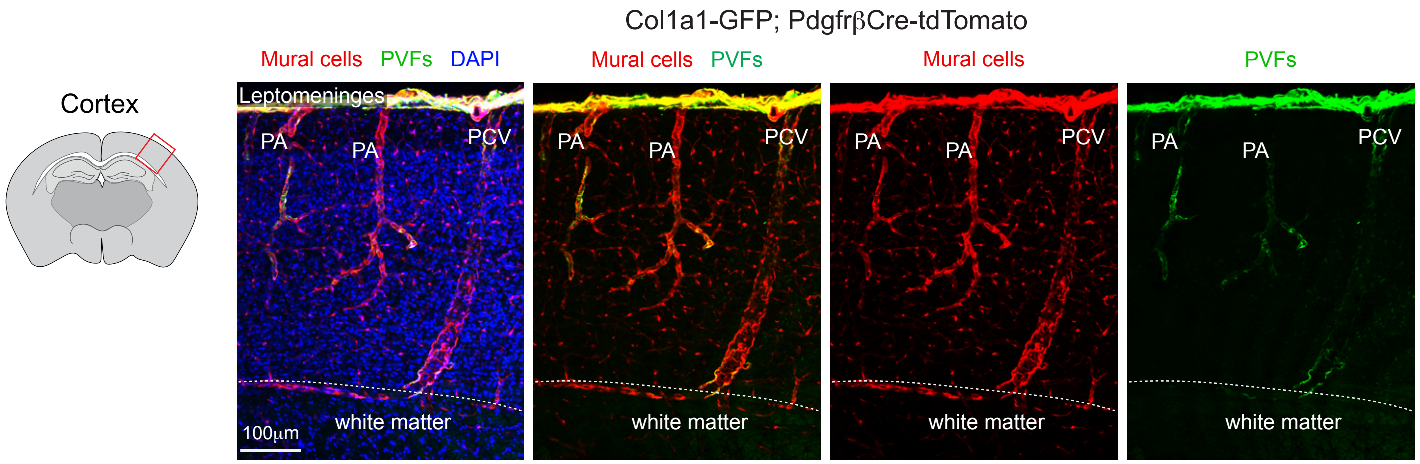


**Supplement Figure 2: Col1a1-expressing perivascular fibroblasts are found on along principle cortical venules that descend from the pia into the white matter.**

Representative confocal image of the cortex from Col1a1-GFP; PdgfrβCre-tdTomato mice showing perivascular fibroblasts (PVFs; green) are found along the entire length of the large principal component venules (PCVs) extending into the white matter. PVFs can also be found along penetrating arterioles (PA) and pre-capillary regions confirming observations from *in vivo* imaging experiments. Mural cells are shown in red and DAPI in blue.

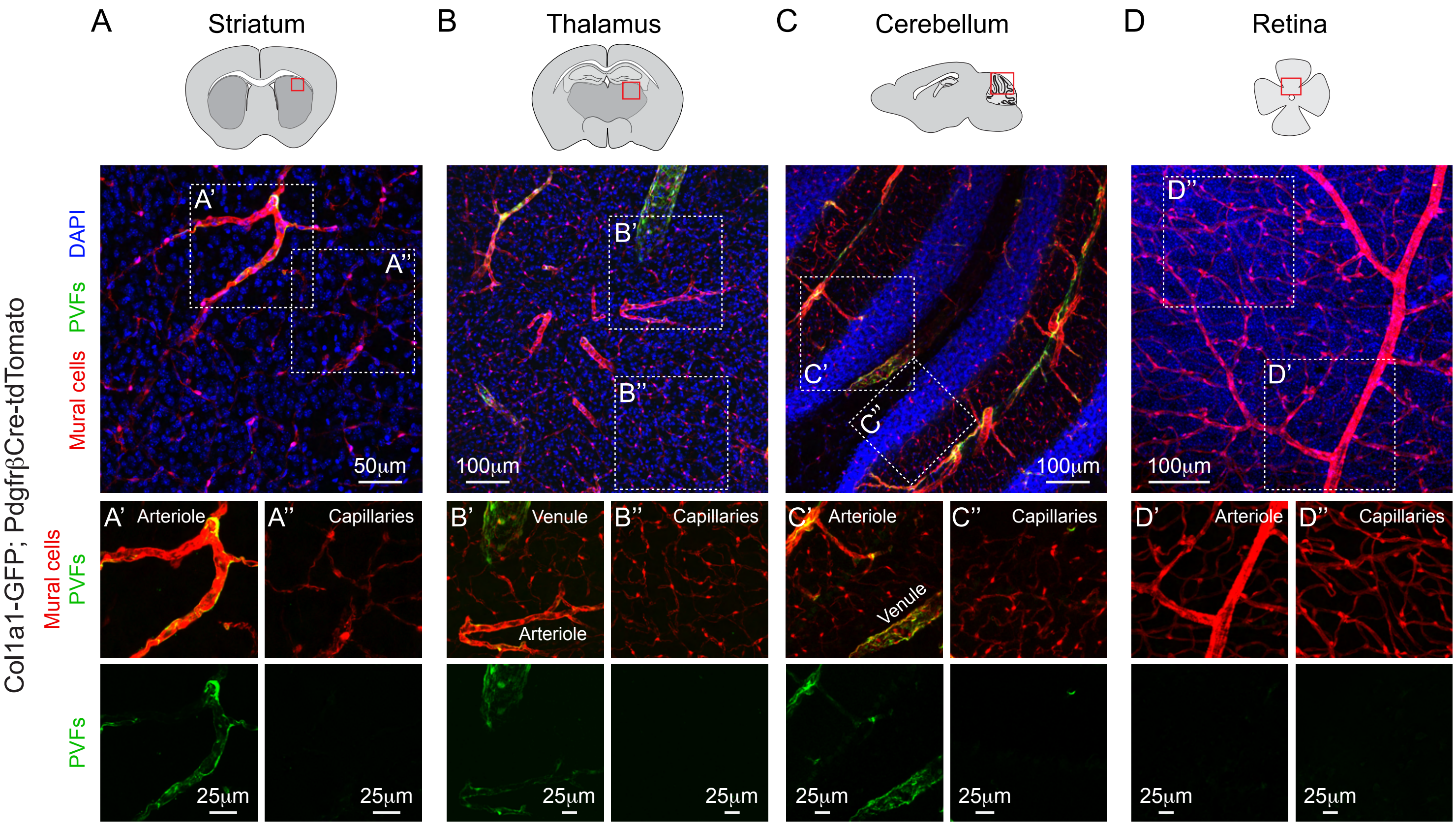


**Supplemental Figure 3: Topological organization of PVFs is similar among different brain regions but PVFs are absent in retina.**

Representative confocal image of the **(A)** striatum, **(B)** thalamus, **(C)** cerebellum and **(D)** retina from Col1a1-GFP; PdgfrβCre-tdTomato mice. Isolated images demonstrate that perivascular fibroblasts (PVFs; green) are found along arterioles, pre-capillary zones and venules in the **(A')** striatum, **(B')** thalamus and **(C')** cerebellum. However, PVFs are not present on the capillary zone as depicted by thin-strand and mesh pericytes (red) in the **(A'')** striatum, **(B'')** thalamus and **(C'')** cerebellum. **(D' and D'')** PVFs are absent on the retinal vasculature. Mural cells are shown in red and DAPI in blue.

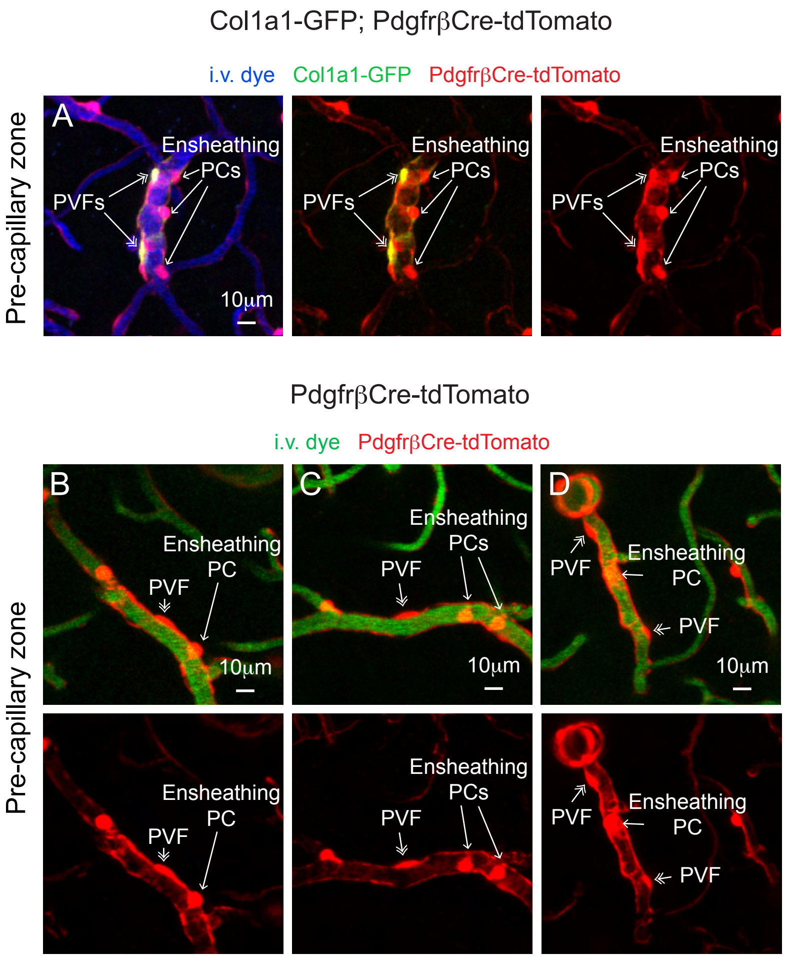


**Supplemental Figure 4: PVFs are morphologically identifiable along the pre-capillary zone and intermingled with ensheathing pericytes**

**(A)** *In vivo* two-photon image of a pre-capillary zone within the somatosensory cortex from Col1a1-GFP; PdgfrβCre-tdTomato mice with ensheathing pericytes (red) and PVFs (green) along pre-capillary vessels (blue; Alexa-680 dextran 2MDa). PVFs are morphologically identifiable due to their flat oblong soma where-as ensheathing have the typical protruding cell body of pericytes.

**(B-D)** *In vivo* two-photon images of three separate pre-capillary zones within the somatosensory cortex from PdgfrβCre-tdTomato mice, arrows point to ensheathing pericytes, double arrows indicate putative PVFs. Vasculature labeled with i.v. administration of FITC-dextran (green; 70kDa).

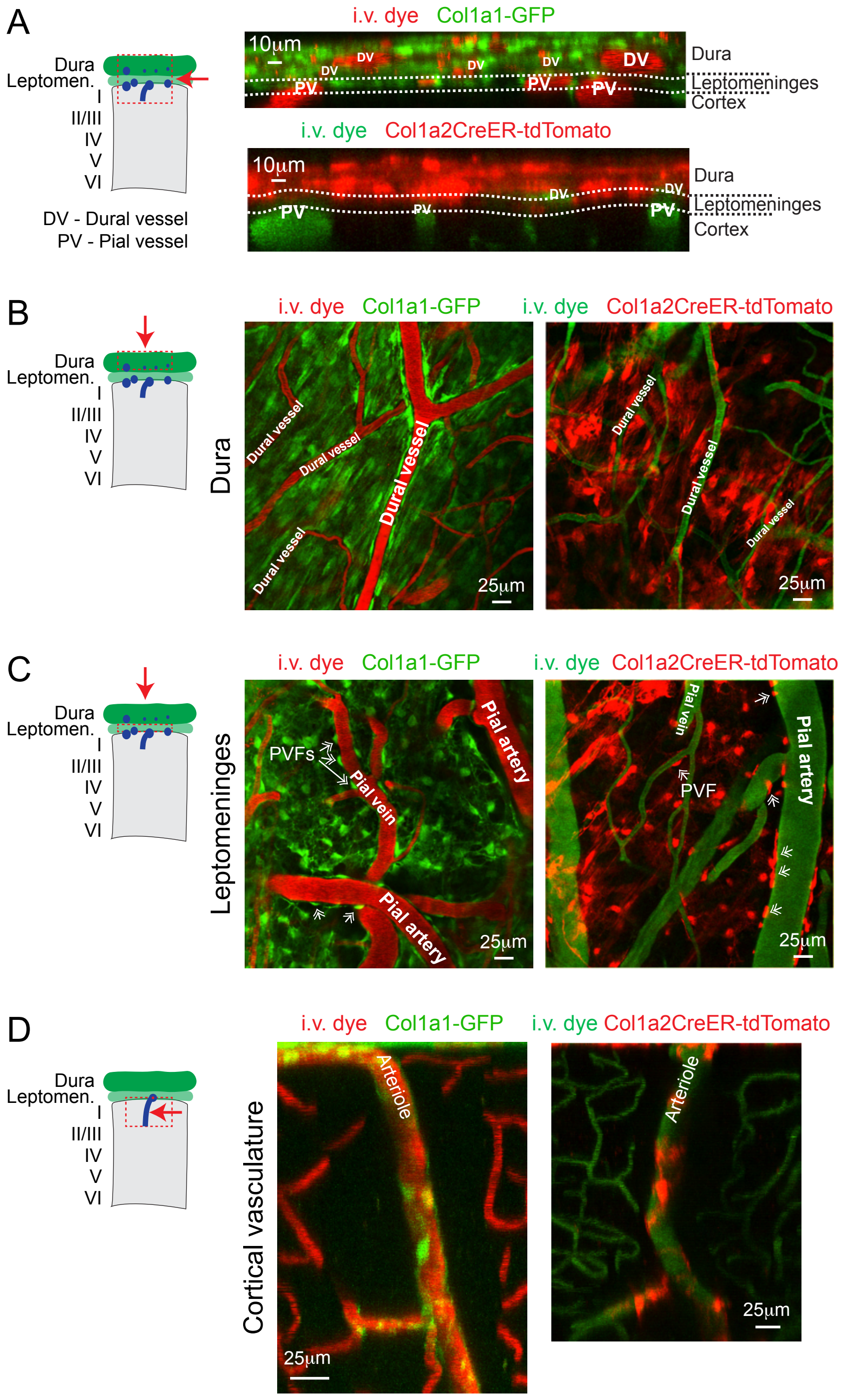

A
Dura
Leptomen.
I
II/III
IV
V
VI
DV - Dural vessel
PV - Pial vessel
i.v. dye Col1a1-GFP
10μm
DV
PV
Dura
Leptomeninges
Cortex
i.v. dye Col1a2CreER-tdTomato
B
Dura
i.v. dye Col1a1-GFP
i.v. dye Col1a2CreER-tdTomato
Dural vessel
25μm
C
Leptomeninges
PVFs
Pial vein
Pial artery
PVF
D
Cortical vasculature
Arteriole

**Supplemental Figure 5: Col1a2CreER mouse line has similar recombination pattern as Col1a1-GFP reporter mice in the adult meninges and cortical vasculature.**

**(A)** Side projections comparing of the meningeal layers (dura and leptomeninges) from *in vivo* two-photon imaging of Col1a1-GFP (top) and Col1a2CreER-tdTomato (bottom) mice showing similar expression patterns within the meninges. Col1a2CreER-tdTomato mice were given 2 consecutive days of tamoxifen (80mg/kg) for sparse labeling experiments. Vasculature labeled with i.v. administration of Texas Red (red; 70kDa) in Col1a1-GFP mice and FITC-dextran (green; 70kDa) in Col1a2CreER-tdTomato mice. Dural vessels (DV) and pial vessels (PV) allowed for the differentiation of the different meningeal layers.
**(B)** Top down, max projection of the dural layers from Col1a1-GFP (left) and Col1a2CreER-tdTomato (right) mice demonstrated similar expression patterns within this meningeal layer.
**(C)** Top down, max projection of leptomeninges (arachnoid and pia) from Col1a1-GFP (left) and Col1a2CreER-tdTomato (right) mice demonstrated similar expression patterns within these meningeal layers. Perivascular fibroblasts (PVFs) (double arrows; GFP+ or tdTomato+) can also be found in both mouse lines.
**(D)** Side projections of the cortical vasculature Col1a1-GFP (left) and Col1a2CreER-tdTomato (right) mice showing similar labeling of PVFs along cortical arterioles.

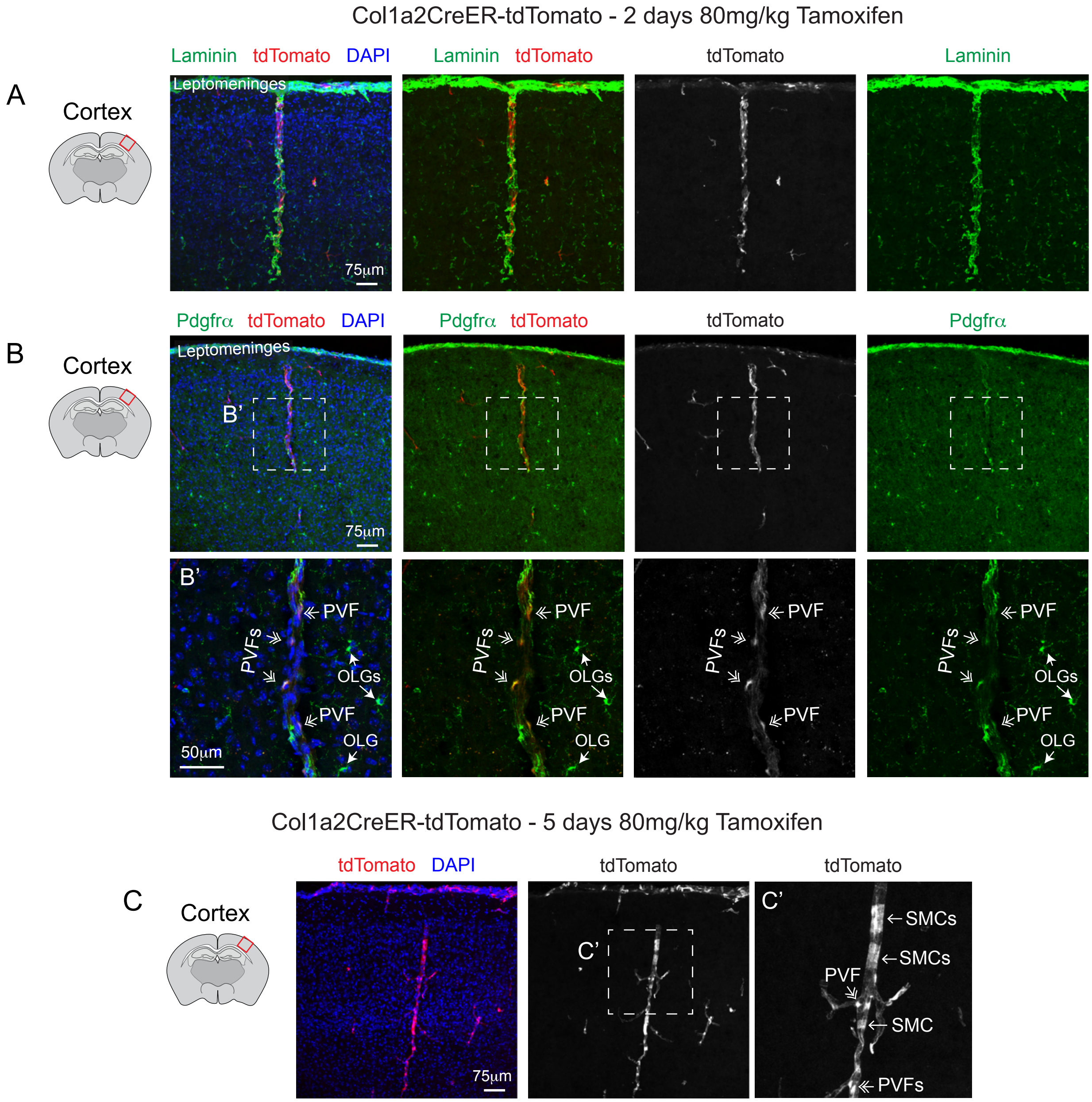


**Supplemental Figure 6: Col1a2CreER recombines in cortical perivascular fibroblasts.**

**(A)** Representative confocal image of the cortex from Col1a2CreER-tdTomato mice treated with 2 consecutive days of tamoxifen (80mg/kg) showing Col1a2-Cre activity recombines in cells (red) along large laminin-labeled vessels (green). DAPI in blue.
**(B)** Immunohistochemistry confirms that tdTomato+ cells along blood vessels express the fibroblast marker Pdgfrα. Double arrows denote putative PVFs. Pdgfrα is also expressed by oligodendrocytes (OLGs).
**(C)** Representative confocal image of the cortex from Col1a2CreER-tdTomato mice treated with 5 consecutive days of tamoxifen (80mg/kg) showing Col1a2-Cre activity recombines in some smooth muscle cells and perivascular fibroblasts.

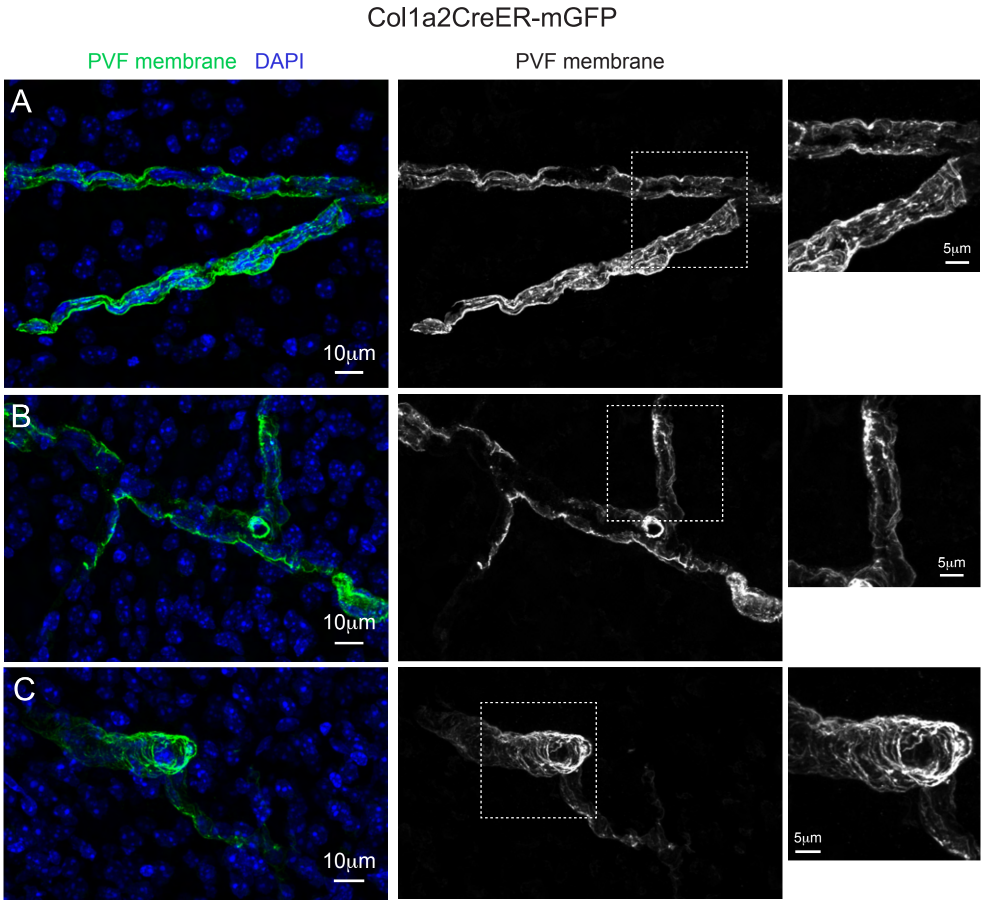


**Supplemental Figure 7: Perivascular fibroblasts have ruffled sheet-like processes.**

**(A-C)** Multiple high-resolution confocal images of PVF membrane (green) from Col1a2CreER-mGFP mice given 2 consecutive days of tamoxifen (80mg/kg). Isolated image demonstrates the intricate ruffled membranes of the PVF lamellae. DAPI staining in blue.

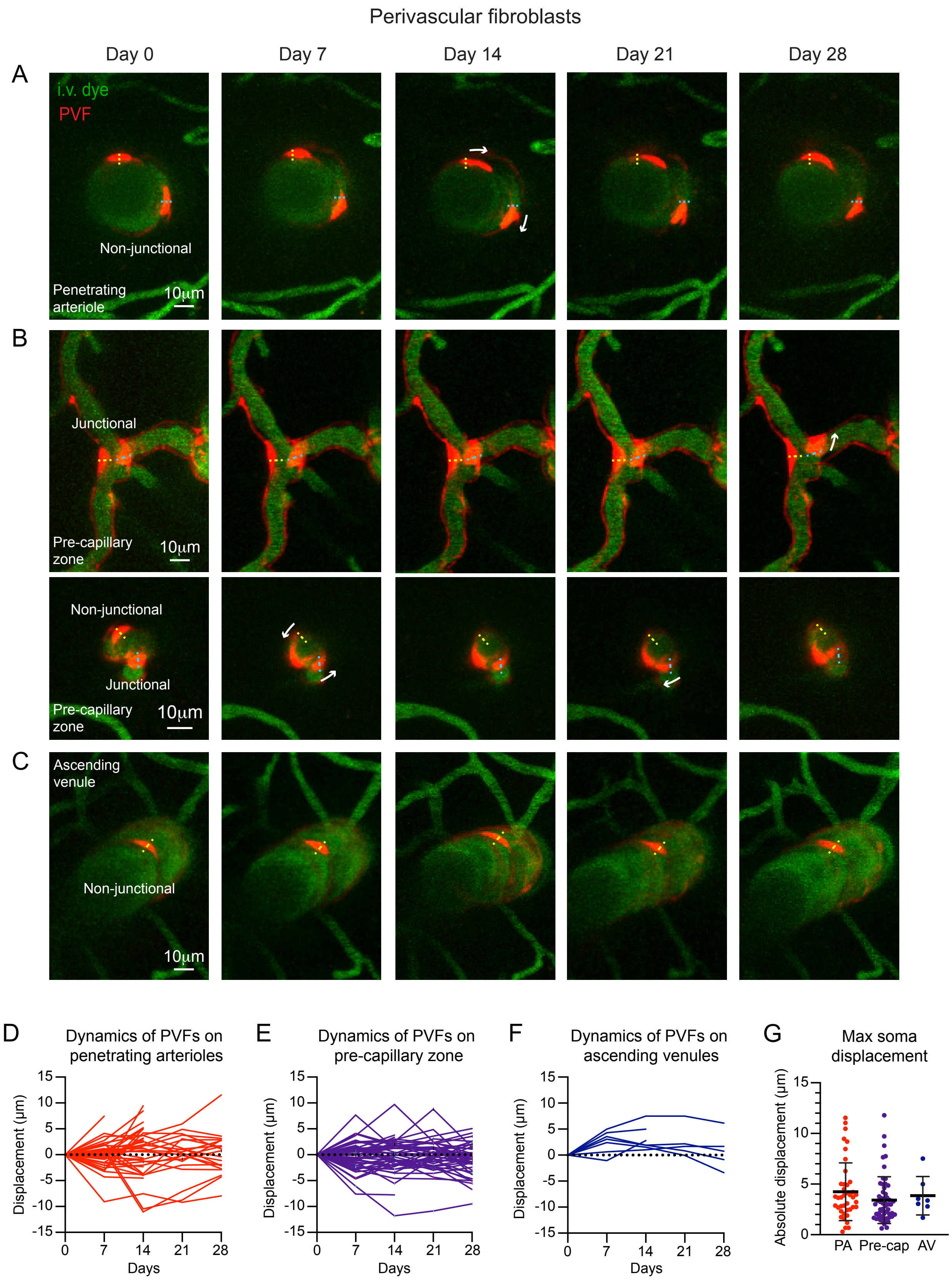

Perivascular fibroblasts
Day 0
Day 7
Day 14
Day 21
Day 28
A
i.v. dye
PVF
Non-junctional
Penetrating arteriole
10μm
B
Junctional
Pre-capillary zone
10μm
Non-junctional
Junctional
Pre-capillary zone
10μm
C
Ascending venule
Non-junctional
10μm
D
Dynamics of PVFs on penetrating arterioles
Displacement (μm)
Days
E
Dynamics of PVFs on pre-capillary zone
Displacement (μm)
Days
F
Dynamics of PVFs on ascending venules
Displacement (μm)
Days
G
Max soma displacement
Absolute displacement (μm)
PA
Pre-cap
AV

**Supplemental Figure 8: Dynamics of perivascular fibroblasts is comparable across different vascular zones.**

**(A-C)** Representative *in vivo* two-photon images of perivascular fibroblasts (PVF; red) over 28 days from Col1a2CreER-tdTomato mice treated with 2 consecutive days of tamoxifen (80mg/kg) along **(A)** penetrating arterioles, **(B)** pre-capillary zones and **(C)** ascending venules. Dashed line indicates initial soma position on Day 0. Vasculature labeled with i.v. administration of FITC-dextran (70kDa; green).
**(D-F)** Graph demonstrating soma displacement of PVFs over 28 days from initial position on day 0 along **(D)** penetrating arterioles (n=41 PVFs), **(E)** pre-capillary zone (n=54) and **(F)** ascending venules (n=7) from 4 Col1a2CreER-tdTomato mice.
**(F)** Graph comparing maximum displacement of PVFs along penetrating arterioles (PA), pre-capillary zone and ascending venule (AV). Analyses demonstrated that dynamics of PVFs across different vascular zones is similar (Kruskal-Wallis test H=3.023, p=0.2205. Mean rank: PA=57.51, Pre-cap=47.22, AV=57.29).
